# NNeurite: artificial neuronal networks for the unsupervised extraction of axonal and dendritic time-lapse signals

**DOI:** 10.1101/2022.01.11.475549

**Authors:** Nicolas Chenouard, Vladimir Kouskoff, Richard W. Tsien, Frédéric Gambino

## Abstract

Fluorescence microscopy of Ca^2+^ transients in small neurites of the behaving mouse provides an unprecedented view of the micrometer-scale mechanisms supporting neuronal communication and computation, and therefore opens the way to understanding their role in cognition. However, the exploitation of this growing and precious experimental data is impeded by the scarcity of methods dedicated to the analysis of images of neurites activity *in vivo*. We present NNeurite, a set of mathematical and computational techniques specialized for the analysis of time-lapse microscopy images of neurite activity in small behaving animals. Starting from noisy and unstable microscopy images containing an unknown number of small neurites, NNeurite simultaneously aligns images, denoises signals and extracts the location and the temporal activity of the sources of Ca^2+^ transients. At the core of NNeurite is a novel artificial neuronal network (NN) which we have specifically designed to solve the non-negative matrix factorization (NMF) problem modeling source separation in fluorescence microscopy images. For the first time, we have embedded non-rigid image alignment in the NMF optimization procedure, hence allowing to stabilize images based on the transient and weak neurite signals. NNeurite processing is free of any human intervention as NN training is unsupervised and the unknown number of Ca^2+^ sources is automatically obtained by the NN-based computation of a low-dimensional representation of time-lapse images. Importantly, the spatial shapes of the sources of Ca^2+^ fluorescence are not constrained in NNeurite, which allowed to automatically extract the micrometer-scale details of dendritic and axonal branches, such dendritic spines and synaptic boutons, in the cortex of behaving mice. We provide NNeurite as a free and open-source library to support the efforts of the community in advancing *in vivo* microscopy of neurite activity.

## 1 Introduction

Optical microscopy of genetically-encoded fluorescent Ca^2+^ sensors offers distinctive advantages for recording a proxy of neural activity in small behaving animals: it gives access to both the activity and topology of large neuronal ensembles [1], genetics allows selective targeting of fluorescent probes to defined neuronal sub-groups based on their genotype [2], and it allows day-to-day tracking of the activity of the same biological targets, over weeks of experiments [3]. *In vivo*, Ca^2+^-imaging was first limited to large cell body compartments which offer superior signal-to-noise-ratio (SNR) [4]. Efficient and accurate processing pipelines have been designed and are now freely available for experimentalists to analyze somatic Ca^2+^ data [5–12]. They allow stabilizing moving microscopy images [13], denoising them [7], extracting the location of cell bodies and their temporal activity [7], as well a deconvolving the Ca^2+^ temporal activity of somas to infer action potential numbers [5, 7, 14, 15]. At the micrometer scale, small dendritic features have been increasingly under focus and *in vivo* microscopy contributed to insights in dendritic signal integration [16] and topological and functional spine plasticity [17–19], which translated in advances in understanding the mechanisms of cognition and perception [20,21]. On the other side of the synapse, imaging the activity of axonal projections from one brain region to another is now emerging as a precious tool for systems neuroscience: it gives access to the specific information communicated between two brain regions of interest [22–25]. Nevertheless, the exploitation of the wealth of experimental data for small neurites is hampered by the scarcity of robust analysis tools. Indeed, the analysis of fluorescent signals from small neurites poses challenges distinct from that of somatic imaging: (1) SNR is markedly lower because of the small number of fluorescent probes in such confined volumes, with often no detectable fluorescence during inactivity, (2) the thin and complex shapes of the fluorescent domains such as dendritic trees and spines or axonal branches do not conform to a predefined shape, (3) imaging these micrometer-scale features *in vivo*, in particular in behaving animals, is critically susceptible to motion artifacts of the same scale.

NNeurite (‘artificial Neuronal Networks for neurites analysis’) is a set of new analysis methods which are dedicated to the analysis of fluorescent images of small neurites. NNeurite relies on a new formulation of the Ca^2+^domain extraction process with an unsupervised artificial neuronal network (NN). Crucially for neurite image analysis, NNeurite integrates in a single optimization procedure signal denoising, image non-rigid stabilization and source shape and temporal activity inference. Moreover, complex micrometer-scale shapes are favored since no morphological constraint is imposed. We tested NNeurite with Ca^2+^ images of both dendrites and axons of behaving mice, but the framework is generic enough to be applied to other types of fluorescent markers of activity. A full implementation of NNeurite as a Python library with graphical processing unit (GPU) based acceleration (Tensorflow implementation) is freely and openly provided^1^ for the community to use and adapt.

## 2 Image model

### 2.1 Non-negative matrix factorization-based source extraction

In the linear regime of fluorescent probes, Ca^2+^ sensors in particular, a simple microscopy image model consists in

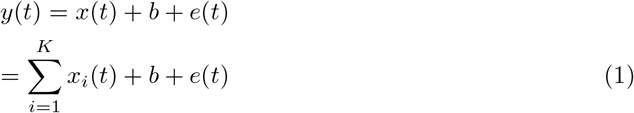

where 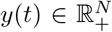 is the column vector of *N* recorded pixel values at time index 1 ≤ *t* ≤ *T*. It corresponds to the combination of the fluorescence contribution of

- a set of *K* Ca^2+^ sources: 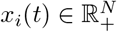 for *i* = 1, …, *K*,
- a fixed background: 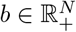,
- some random noise: *e*(*t*) ∈ ℝ*^N^*.

Here, we use the term ‘source’ for a spatial domain with synchronized changes of Ca^2+^ concentration, such as an axonal or dendritic segment. Still, fluorescence intensity in a single domain can be heterogeneous in space due to local variations in Ca^2+^ concentration, geometry of the domain and the availability of the fluorescent probe. These considerations are well-matched by a multiplicative model between a spatially changing kernel 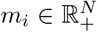, often called ‘footprint’, and a representative ‘amplitude’ 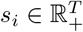 modeling time variations [7, 26]:

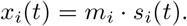

Inserting this expression in (1), we compactly rewrite the image model as:

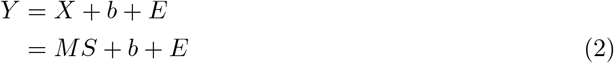

where matrices conform to: 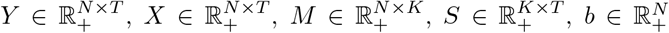 and 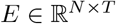.

For microscopy data analysis, the crux of the problem is to decipher the product *MS* with *M* ≥ 0 and *S* ≥ 0, which corresponds to a non-negative matrix factorization (NMF) problem [7, 26]. The general NMF problem takes the form:

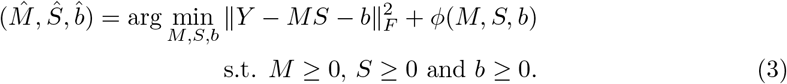

The real-valued function *φ*, called ‘regularizer’, is used to guide optimization towards desirable outcomes. Indeed, NMF is a non-convex, NP-hard problem and heuristic methods may struggle to avoid local minima and initialization-dependence without proper regularization. The *ℓ*_1_-norm (||*u*_1_|| = Σ_*k*_ |*u_k_*|) is a typical regularization function which can promote sparsity of the neural activity or of the footprints [5, 7, 9]. Implicit forms of regularization have also been used under the form of heuristic rules during footprint and activity estimation (e.g. connected non-zero pixels for footprints [6, 7]). However, strong regularization may come at the cost of simplistic solutions which cannot approximate with high accuracy the complex features of neurite microscopy imaging, such as the complex shapes of the Ca^2+^ domains.

### 2.2 Spatially unstable time-lapse images

Time-lapse imaging microscopy is generally corrupted by slow spatial drifts of the sample which invalidate the static image model (Eq 1). This issue is exacerbated for *in vivo* microscopy in behaving animals, despite head-fixation, because motor actions can produce deformations of the field of view several times the size of neurites at the sub-second scale. A standard procedure to minimize motion artifacts for Ca^2+^ image analysis has been to globally, or locally, align images prior to source-extraction in an attempt to conform to the static image model compatible with NMF analysis [8, 11]. In this case, image registration is classically performed with respect to a slowly-changing template of the images [13], which can be ill-suited to neurite image processing for which no stable landmark of fluorescence exist because of the low, or non-existent, fluorescence at rest. To cope with this issue, we propose that a modified image model which complies with rapid field-of-view deformations is:

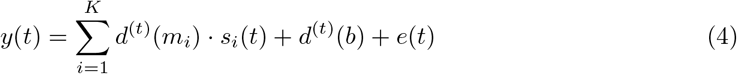

where *d*^(*t*)^ denotes a non-rigid spatial deformation of the field-of-view at time point *t*. We therefore rewrite the NMF-based source-extraction problem as:

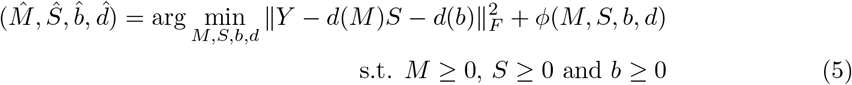

the set of deformations *d* = {*d*^(*t*)^}_*t*=1,…,*T*_ can also be regularized. We argue that optimizing this combined formulation would allow using the transient fluorescence of Ca^2+^ sources to robustly align images as they are identified. However, no computational method has been proposed so far to directly solve this upgraded problem.

## 3 Proposed methods: simultaneous image registration and NMF-based source extraction with artificial NNs

### 3.1 NN-based NMF of microscopy images

#### 3.1.1 Principles

In order to solve the NMF problem (Eq 3) we propose a new NN architecture (Fig 1) with distinctive features. It has two main parts: 1) a sequential set of layers which project each individual image *y*(*t*) to a, low, *K*_0_-dimensional space and 2) a linear layer which synthesizes images *z*(*t*) from their low-dimensional representation. A key concept underlying our approach is that for image data conforming to the NMF model (Eq 2) there must exist a function 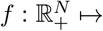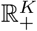 which can extract a good approximation of the amplitude of the *K* sources for each input image *y*(*t*). Intuitively, for footprints which do not overlap, one typical such function would average the measured pixel values at the location of each footprint and correct for noise and background components. For overlapping footprints, extra corrections would need to be applied to separate fluorescence contributions from different sources. In the proposed NN framework, the image model (Eq 2) becomes:

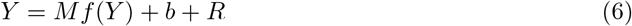

where *f* (*Y*) = [*f* (*y*(1)), …, *f* (*y*(*T*))] (*source amplitude extraction*) and *Mf* (*Y*) (*image synthesis*) are implemented in the left and right parts of the NN in Fig 1. We propose to deviate from the standard NMF formulation (Eq 3) and carry out parameter optimization over the footprints *M*, the amplitude extraction function *f* (instead of the amplitudes *S*) and the background *b* as:

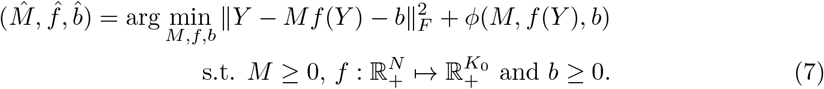

**Figure 1:**
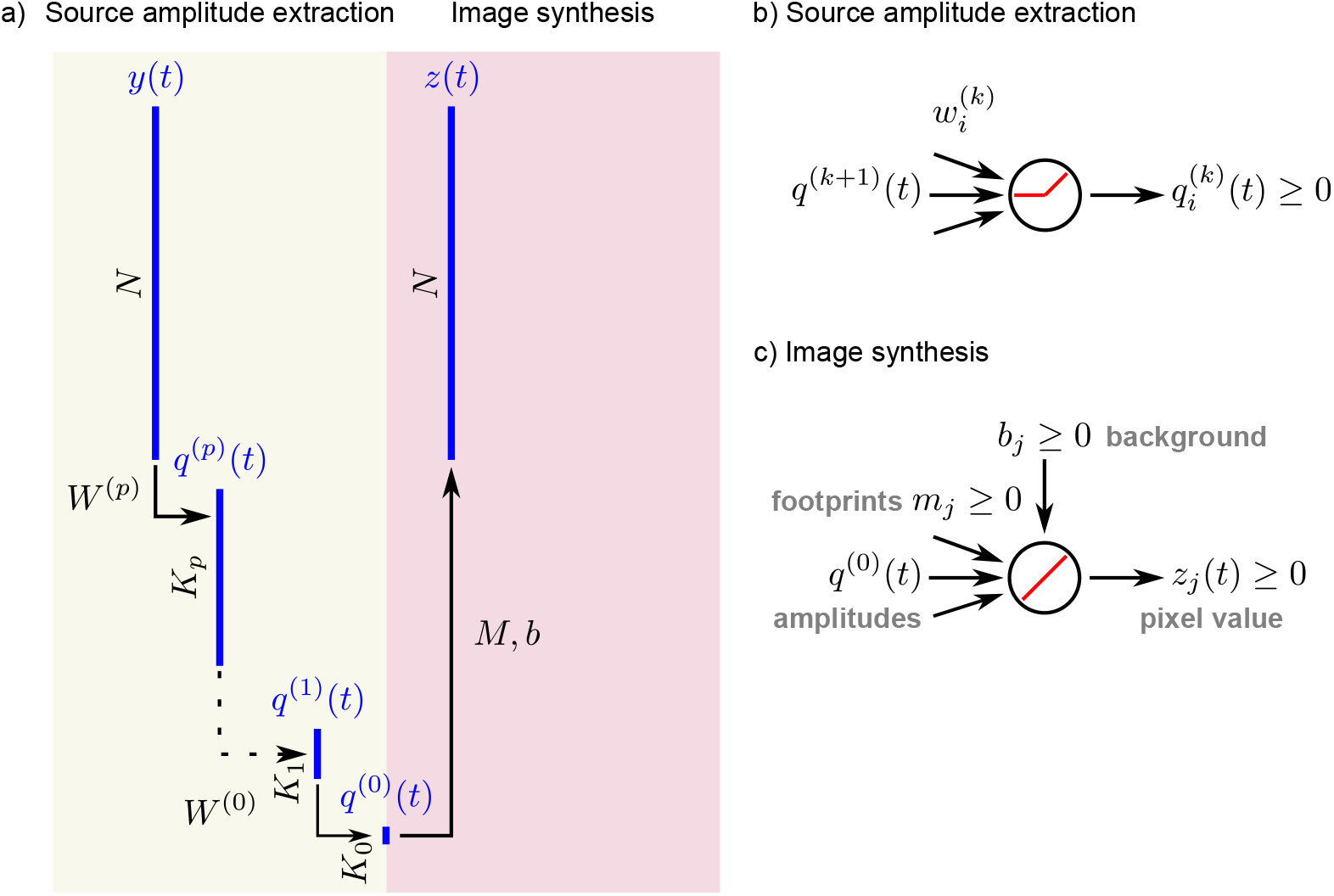
Artificial neuronal network for identifying Ca^2+^ sources in neuronal microscopy images. a) The main architecture of the proposed NN for NMF is composed of two main parts. First, a compressive series of layers is applied (yellow), ultimately computing a low (*K*_0_)-dimensional representation (*q*^(0)^(*t*)) which estimates the source amplitudes 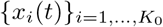 for a single image frame *y*(*t*). Then, an expansive synthesis part based on a single layer (red) computes the image approximation *z*(*t*). b) Neuronal units of the amplitude extraction path are densely connected, with weights 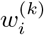 at level *k* for the *i*-th unit, and use an unbiased ReLU activation function [28] to produce positive coefficients 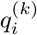. c) Image synthesis is based on a single layer of units with linear integration to match the linear image model (Eq 2). One unit is used for each pixel, with the bias value *b_j_* corresponding to the background at the *j*-th pixel, and the coefficients {*m_j_*}_*j*=1*,…,K*_0 as the source footprints values at this pixel.

One could also view the proposed NN as an *autoencoder* for dimensionality reduction [27], but with a decoding (synthesis) part of the network which strictly matches the fluorescence image model.

#### 3.1.2 Amplitude extraction as a low-dimensional image projection

The feasible space for *f* is determined by the inter-layer connectivity, the type of activation function, and the depth of the network. In particular, deeper NNs with non-linear activation functions allow for more complex calculations. We chose densely-connected layers with unbiased rectified linear unit (ReLU) [28] activation :

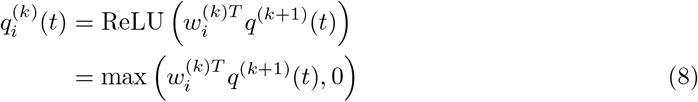

for the *i*-th unit at level *k*, or

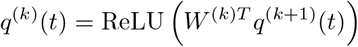

where 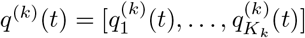 is the grouped decomposition of *y*(*t*) at level *k* (Fig 1). Here, 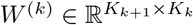 is the matrix of parameters for input integration of the neuron unit.

#### 3.1.3 NN-based image synthesis

Image synthesis is linear with respect to *f* (*Y*) in Eq 7. After source amplitude extraction, we therefore implement synthesis as a single NN layer with a linear, biased activation function (Fig 1a, c):

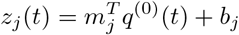

for *j* = 1, …, *N*. 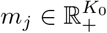 is the vector of footprint values at the *j*th pixel for the *K*_0_ sources. *b_j_* ≥ 0 is the bias of the activation function for the *j*th neuronal unit and models the background contribution to measurements. Pooling together the *N* pixels, we can write:

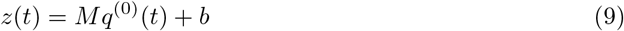

with 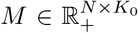 and 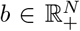 for the synthesis layer. The synthesis equation (Eq 9) matches in all aspects the image model (Eq 2), except for the absence of the noise component. The proposed NN network (Eq 1) can therefore be viewed as as de-noising algorithm working through structured, low-(*K*_0_)-rank approximation of images.

#### 3.1.4 NN training

By inserting Eq 9 in the minimization problem (Eq 7) without regularization we obtain a NN parameter optimization problem with *ℓ*_2_ loss:

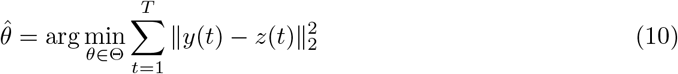

where *z*(*t*) is the output of the NN with parameters *θ* = *W*^0^, …, {*W^p^, M, b*}. Here, the feasible parameter space Θ is limited only by the non-negativity constraints of the image model: *M* ≥ 0 and *b* ≥ 0. Source amplitude non-negativity (*q*^(0)^(*t*) ≥ 0) is warranted by the ReLU activation function for source amplitude extraction (Eq 8). The minimization problem (Eq 10) has a standard form and can therefore be solved by using stochastic gradient descent (SGD)-type methods for NN optimization. NMF regularization should contribute extra terms to error gradient computation in SGD. Interestingly, the most common regularization terms – *ℓ*_1_ or *ℓ*_2_ penalization of source activity or footprints – directly translate to regularization terms in the NN framework:

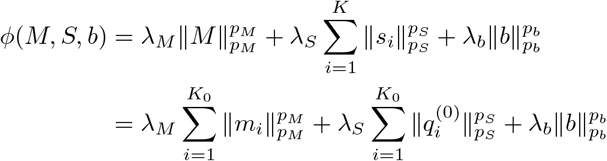

with *p_Y_, p_M_, p_b_* ∈ {1, 2}, and *λ_S_, λ_M_, λ_b_* ∈ ℝ_+_. Most implementations of SGD algorithms for NNs readily handle such regularization terms because they are commonly used to prevent data over-fitting in supervised learning tasks.

An important advantage of the proposed problem (Eq 7) is that optimization does not explicitly aim at estimating the temporal amplitudes for all the image frames (*s_i_*(*t*) for *t* = 1, …, *T*, as in Eq 5), but instead at computing the NN parameters which can be used to estimate them on the fly with a series of inexpensive calculations (*f* (*y*(*t*)), the analysis NN layers filtering). In order to markedly speedup training of the NN we therefore propose to carry out optimization over a small subset of chosen image frames Ω before extracting source amplitudes over the whole set of available frames:

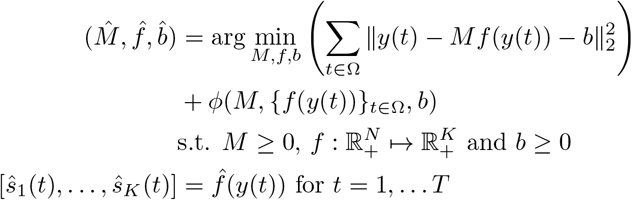

In practice, since dendritic and axonal signals are often temporally sparse in the cortex we can select a small subset of frames in which sources are active and leave out most inactivity frames. In a first approach our strategy was to use the highest-intensity image frames (*t*s with highest ||*y*(*t*)||_2_ for 1 ≤ *t* ≤ *T*) mixed-in with a small number of random images with low fluorescence intensity (*t*s with small ||*y*(*t*)|| _2_) to account for source inactivation.

#### 3.1.5 NMF regularization for low-dimensional signal coding

A low-dimensional representation of time-lapse images can be obtained by design of the NN architecture: choosing a low value *K*_0_ for the *a priori* maximum number of sources as a good approximation of the ‘true’ number of sources, and optimizing the NN parameters for the problem (Eq 11). Instead we propose a more flexible approach which we found to rely less on prior knowledge: regularization of the source footprints and amplitudes. Doing so, naive NN networks start by approximating images with high (*K*_0_)-dimensional models and progressively reduce the number of required sources through parameter optimization. Instead of standard *ℓ*_1_ or *ℓ*_2_ penalization terms, we propose to adapt group-lasso regularization [29] for our image NMF problem:

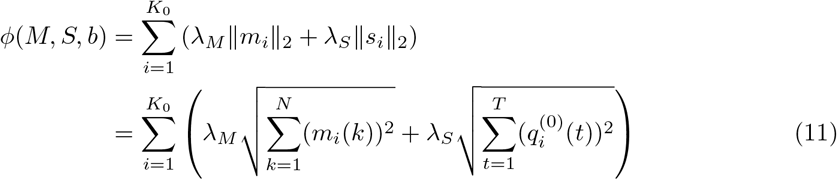

The squared terms in the proposed regularizer ensure that footprints and amplitudes can include low values, allowing sources to span large parts of the image for example, without being over-penalized. Taking the square root of the sum of those squared values and summing over all sources favors models with only few of those large-span footprints and non-zero temporal activity. Importantly, the regularization functions are convex and therefore well-suited to gradient-based numerical optimization. For neurites imaging, the pixel-wise photon-count is low and can typically be modeled as a Poisson process which variance scales with its mean. We therefore tie the regularization strength to the pixel variance: *λ_M_, λ_S_* ∝ var(*Y*). Also, because neurites can show complex topology and activity patterns, we propose to not constrain further the source footprints and temporal amplitudes, and to keep the background term unconstrained.

### 3.2 Combined local NMF procedures

In microscopy recordings of neurite activity, it is typical for a single field of view to comprise tens of sources of fluorescence (eg [22, 25, 30]). In such conditions, solving the large-scale NMF problem for source extraction generally lacks robustness: results may show variability with respect to initialization and parameters, and small changes in the input image data can translate to significant changes in the output. To tackle this issue we propose to define *P* local NMF problems of low (*K′ ≪ K*) dimension:

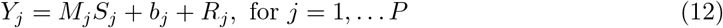

where *Y_j_* is a small image patch. The small size of the local NMF problems allows one to solve them robustly with the proposed NN framework, but also to process them in parallel with distributed computational resources to speed up computation [11]. Their results are gathered in a global model of images where sources spanning multiple NMF problems are combined. As illustrated in Fig 2, these principles map into a NN architecture with locally connected neuronal units at the first layer of the image analysis pipeline and at the last layer of the image synthesis part. The activity of the *K* global sources is calculated as a specific linear combination of the outputs of the local NMF solvers:

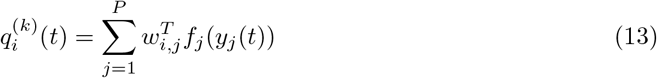

with 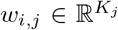, for *i* = 1, … *K*. Here, each local source should contribute exactly one global source. This fits nicely with the framework on non-supervised, hard, classification where each cluster of local sources corresponds to one source at the whole image level. We propose to take advantage of an algorithm based on affinity propagation with an automatically-determined number of clusters [31], where we chose the temporal Pearson’s correlation between the local source amplitudes as their affinity index. We then compute the merging weights *w_i,j_* in each cluster of sources as the first eigenvector of the matrix of temporal amplitudes to minimize the loss of accuracy due to dimensionality reduction.

**Figure 2:**
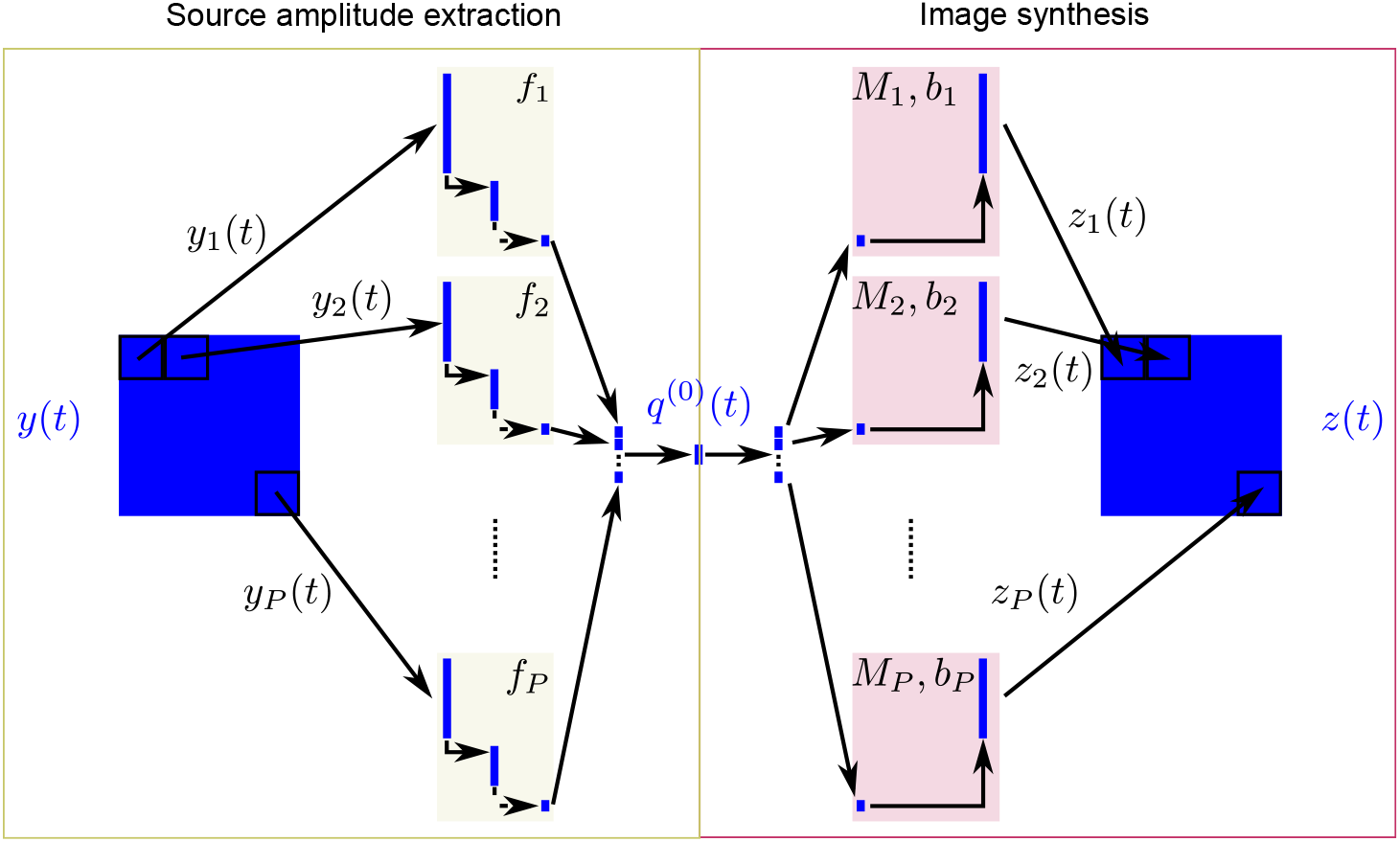
Combined local NNs for distributed NMF. Input images are divided into patches {*y_i_*}_*i=1,…,P*_ and independent local NNs (Fig 1) are trained to obtain the optimized solutions {*f_i_, M_i_, b_i_*}_*i*=1,…,*P*_. The local source amplitude extraction paths (yellow, {*f_i_*}_*i*=1,…,*P*_) are then linearly combined to compute low-dimensional representations *q*^(0)^(*t*) of whole images. Here, *q*^(0)^(*t*) corresponds to the global sources amplitudes at frame *t* and it can be processed (red) to synthesize an approximation of *y*(*t*) by using the local image synthesis functions {*M_i_, b_i_*}_*i*_=1,…,*_P_*.

### 3.3 Simultaneous non-rigid image registration and NMF

When stable landmarks of fluorescence cannot be used to align moving images, we propose that analyzing the transient sources signals is a viable strategy. However, since the location of fluorescent sources is unknown *a priori*, it requires identifying sources and registering images simultaneously (Eq 5). We consider here the useful case of non-rigid transformations which can be approximated by local translations of parts of the image [13] with integer pixel translations *d_δx,δy_* = translation_(*δx,δy*)_(·), where *δx, δy* ∈ ℤ. A crucial advantage of the NNeurite framework is that the analysis layers (*f_i_* functions) can be applied to translated versions of input image tiles (*d_δx,δy_*(*y_i_*)) with a single modification of the first layer: shifting the weights *W^p^* in space to compensate for the translation of the input image (*d_δx,δy_*(*W^p^*)). When all shifts are considered up to a maximal distance, the linear analysis with all *d_δx,δy_*(*W^p^*) weights amounts to spatially convolving the image patches with *W^p^*. Hence, the first dense analysis layer of the NN structure becomes a spatial convolution layer with the same weights *W^p^*. The resulting multi-channel (one per translation) coefficients are then analyzed without modification of the NN network (Fig 3).

**Figure 3:**
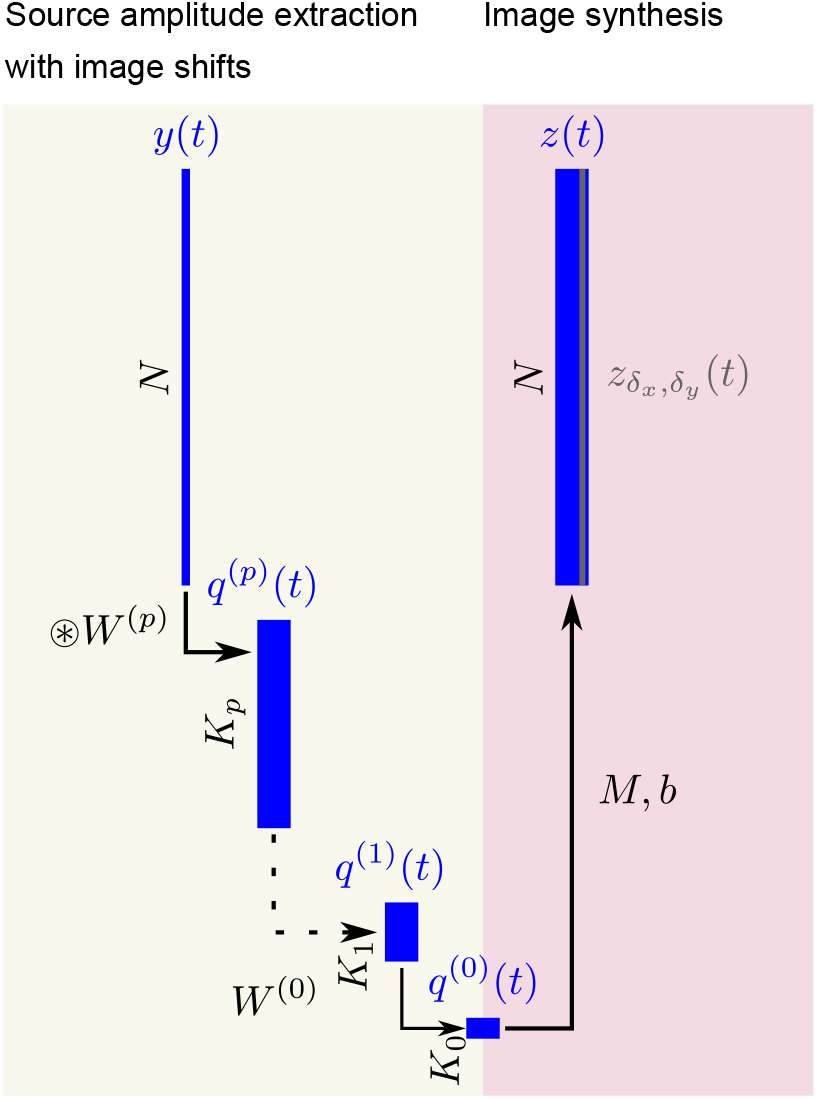
Multi-channel convolutive NN for simultaneous image registration and source extraction. The first analysis layer is a convolution with the dense kernel *W* ^(*p*)^ with unit vertical and horizontal strides, instead of a single densely connected kernel (Fig 1), hence coping for integer spatial shifts (*δ_x_, δ_y_*) of the input image. It produces multi-channel (one for each spatial shift) coefficients which are passed through the whole amplitude extraction and image synthesis layers to produce a set of non-shifted image approximations 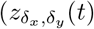, one per shift).

In terms of network optimization, for each local NMF problem, we propose the following cost-function

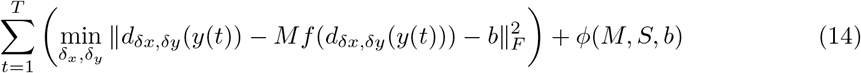

where each channel of the convolutive NN (Fig 3) predicts 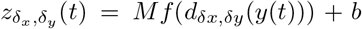 for a different shift (*δ_x_, δ_y_*) of the raw image *y*(*t*). Importantly, our strategy to embed image registration in the NMF problem does not add extra parameters to the original NN for stable images (Fig 1). Therefore, the extra temporal and complexity costs of minimizing Eq 14 for (*M, S, b*) instead of optimizing the original problem (Eq 7) are limited.

### 3.4 Non-regularized source refinement

Posterior to local source grouping into larger ones, we propose to refine source footprints and amplitudes using an alternating optimization strategy:

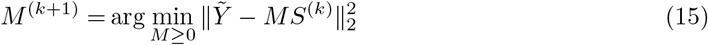

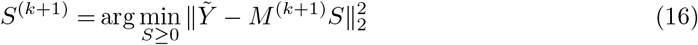

for *k* = 0 to *k*_max_, the maximal number of iterations. Here, 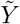 denotes the fluorescent images which have been locally stabilized by compensating for the local translations obtained when optimizing Eq 14. The intuitive strategy of alternating between (15) and (16) to solve NMF is generally impractical without strong regularization because of: 1) its high probability of converging to one of the many local minima of the NMF problem, in the vicinity of the starting point *S*^(0)^, and 2) the unknown numer of sources is fixed *a priori* by the size of *S*^(0)^. We turn these properties into advantages by setting *S*^(0)^ as the amplitudes of the global sources computed by our distributed NNs for NMF (Eq 13), ensuring that the algorithm will refine the output of the NN model without overly diverging from it in terms of source amplitudes, footprints, or number. Therefore, we take advantage of the proposed NN architecture in two ways: performing non-rigid image registration based on neurite fluorescence only, and providing a high-quality input to the unconstrained optimization framework (Eq 15 and 16)

Both (15) and (16) correspond to smooth optimization problems over convex domains. Therefore, they do not admit suboptimal local minima and we can use iterative optimization algorithms to reach global minima. We implemented such optimization procedures by using the framework of the *fast iterative shrinkage-thresholding algorithm* [32].

## 4 Results: processing *in vivo* Ca^2+^signals of neurites

NNeurite accuracy was validated by using a set of *in vivo*, awake, microscopy data generated for this purpose (Fig 4) (Material and methods).

**Figure 4:**
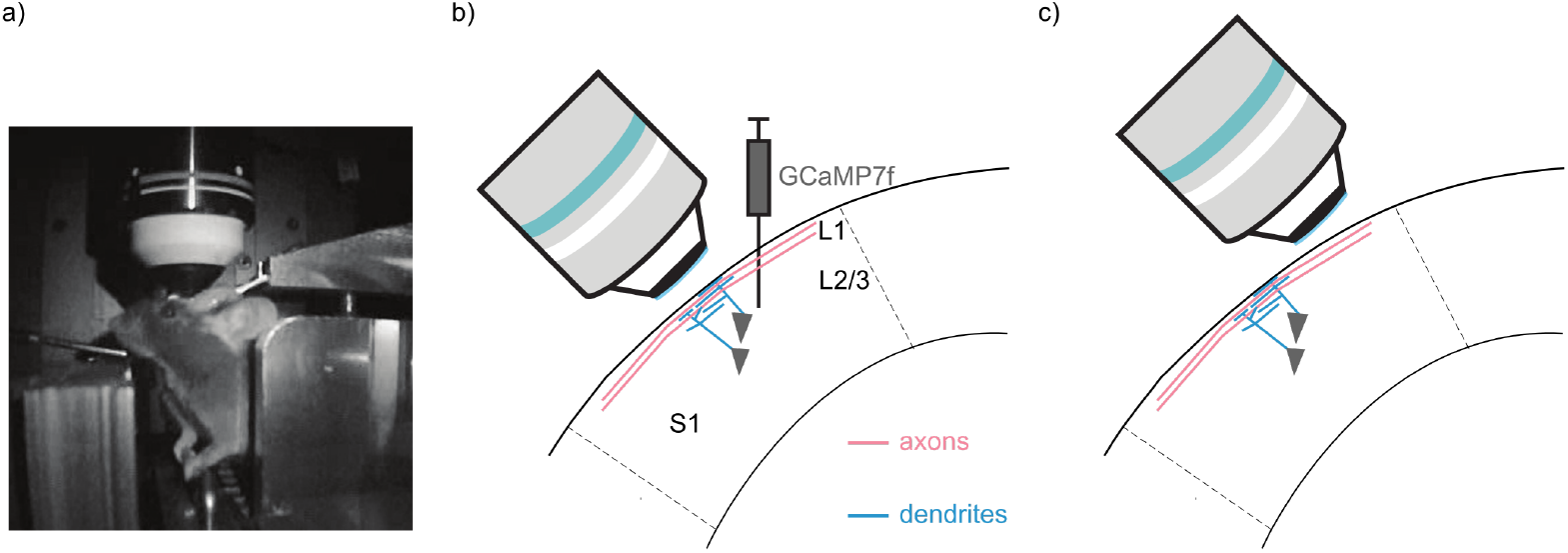
Imaging protocol for axons and dendrites in behaving mice. a) Awake animals were habituated to head-fixation under the microscope. b) Viral injection for expressing the Ca^2+^ sensor GCaMP7f [44] in L2/3 of S1. Microscopy imaging was performed right above the injection point to mostly capture superficial dendritic activity in L1. c) When the microscope objective was moved away from the injection point only far-spreading axonal projections remained [33,34].

### 4.1 Analysis of dendritic signals

When the field-of-view was focused above the virus injection point in the primary somatosensory cortex (S1) at approximately 50-100 *μ*m below the brain surface (Fig 4b), we observed a dense mixture of elongated, spiny and branched structures (Fig 5a) which are typical of dendritic trees of layer (L)2/3 pyramidal cells extending towards the surface and spreading in L1 [33, 34]. Their fluorescent signals were sparse in time and interspersed with out-of-focus short periods when the animals were intensely active. Notably, the high density of dendritic features out of the plane of focus produced a hazy fluorescent background which was also fluctuating. We analyzed 2 min-long movies acquired at 37 Hz with NNeurite. Prior to image processing, out-of-focus images were discarded (Material and methods) and remaining images were re-sampled at 12.3 Hz to enhance the signal-to-noise ratio. The top 20% high intensity patch frames were used for training, with an additional 5% random low-intensity frames. NNeurite was set up to use a maximum of 8 local sources for each 41 × 41 *μ*m rectangular image tile. No specialized image stabilization procedure was applied prior to NNeurite as simultaneous image registration and NMF were performed (Eq 14).

**Figure 5:**
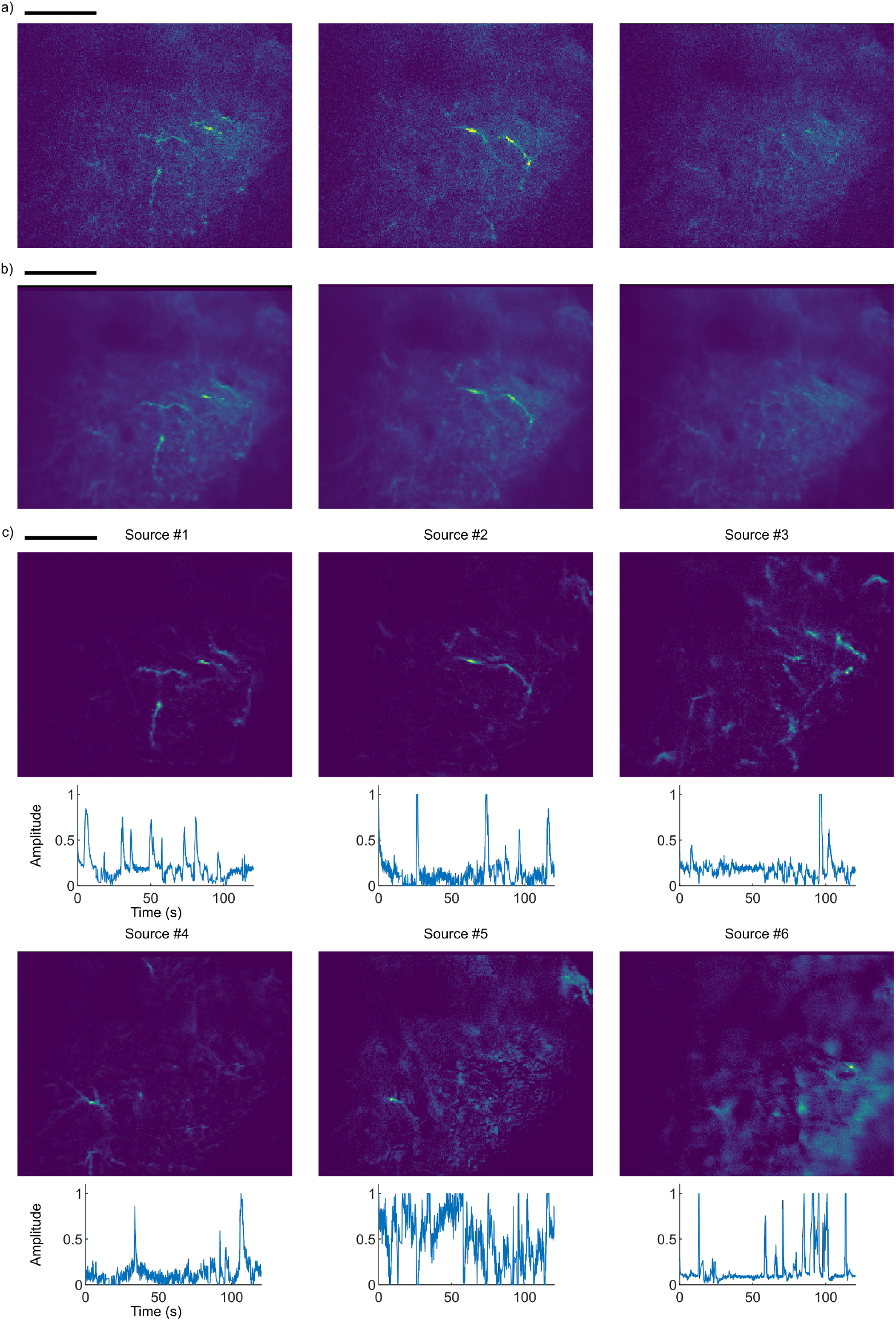
NNeurite extracts dendritic signals in awake behaving mice. a) Exemplar two-photon microscopy images showing transient activation of dendritic features and strong neuropil background. b) Same image frames as (a) analyzed by NNeurite and reconstructed as a noiseless spatio-temporal mixture of sources (*MS* + *b*, Eq 2). c) Example of Ca^2+^ sources extracted by NNeurite for a 2 min image stack in the area. 2S1cale bars: 50 *μ*m.

For an exemplar image sequence, the spatio-temporal mixture model (Eq 2) that was optimized by NNeurite (62 min of processing for 500 iterations of SGD for all 42 patches) provided an accurate approximation of the measured fluorescence (Fig 5b). Both thin dendritic branches with transient short activation and the unstable background neuropil signals were reconstructed, with little tiling effects. Pixel-wise noise was eliminated thanks to its incompatibility with low-dimensional compression, hence showing that, in effects, NNeurite can function as a denoising procedure. For this movie, 86 local sources of fluorescence were identified by the distributed NMF procedure, which were then merged into 10 global sources, 6 of which are shown in Fig 5c. Thanks to the lack of morphological constraint, the footprints of Ca^2+^ sources could assume complex shapes combining, sometimes disjointed, elongated segments (sources #1 to 4), as expected if dendritic trees were ‘cut-across’ by the optical focal plane. Importantly, the spatial resolution of the footprints was at the sub-micrometer scale as small protrusions out of main dendritic shafts, reminiscent of dendritic spines, could be distinguished in some instances (sources #1 to 3). Most amplitude series are near-zero with sparse upticks (sources #1 to 4), in agreement with sparse activation in S1 [35], while some areas with more sustained fluorescence could correspond to autofluorescence artefacts (source #5). Importantly, the lack of morphological constraint for footprints and the ability for them to overlap made it possible to extract out-of-focus neuropil signals as Ca^2+^ sources with delocalized hazy footprints (source #6 and others). The lack of strong correlation between the amplitudes of the widespread neuropil and other sources highlights the ability to unmix the fluorescence of pixels where they overlap.

### 4.2 Analysis of axonal signals

When the field-of-view was moved away (> 300 *μ*m) from the virus injection point (Fig 4c) the fluorescent features became sparser, smoother and thinner (Fig 6a). This description matches that of axons of *L*2*/*3 pyramidal cells which spread laterally in L1 at a greater distance than their dendritic trees [33, 34]. Again, fluorescent signals were sparse in time, subjected to loss-of-focus events, weaker that dendritic transients overall, but the background was less pronounced thanks to the absence of neuropil underneath. Using the same setup of NNeurite as for dendritic image processing, with only modified regularization strength (Material and methods), we again were able to reconstruct accurate representations of the original fluorescence images (Fig 6b) (54 min of processing with 500 iterations of SGD for all 48 patches). Even though noise was mostly eliminated from the reconstruction, NNeurite processing kept intact the complex axonal branches with interspersed swellings – the presumed synaptic boutons. For this 2 min-long movie, 10 local sources of fluorescence were identified were merged into 3 global sources which are shown in Fig 6c. Each source captured a different axonal segment with sparse, asynchronous Ca^2+^ activity (Fig 6c, bottom). The lack of morphological constraint allowed footprints to capture long segments (>50 *μ*m), sometimes branching and ‘zigzagging’ across the field of view (top). For all of them, putative synaptic boutons were clearly identifiable as bright swelling along the thin, elongated filaments. We did not observe spatially-shifted duplicates of the same axonal branches across footprints, which could be caused by transient motion of the field of view during awake imaging, hence proving the ability of NNeurite to effectively stabilize images simultaneously to identifying the axonal sources of fluorescence.

**Figure 6:**
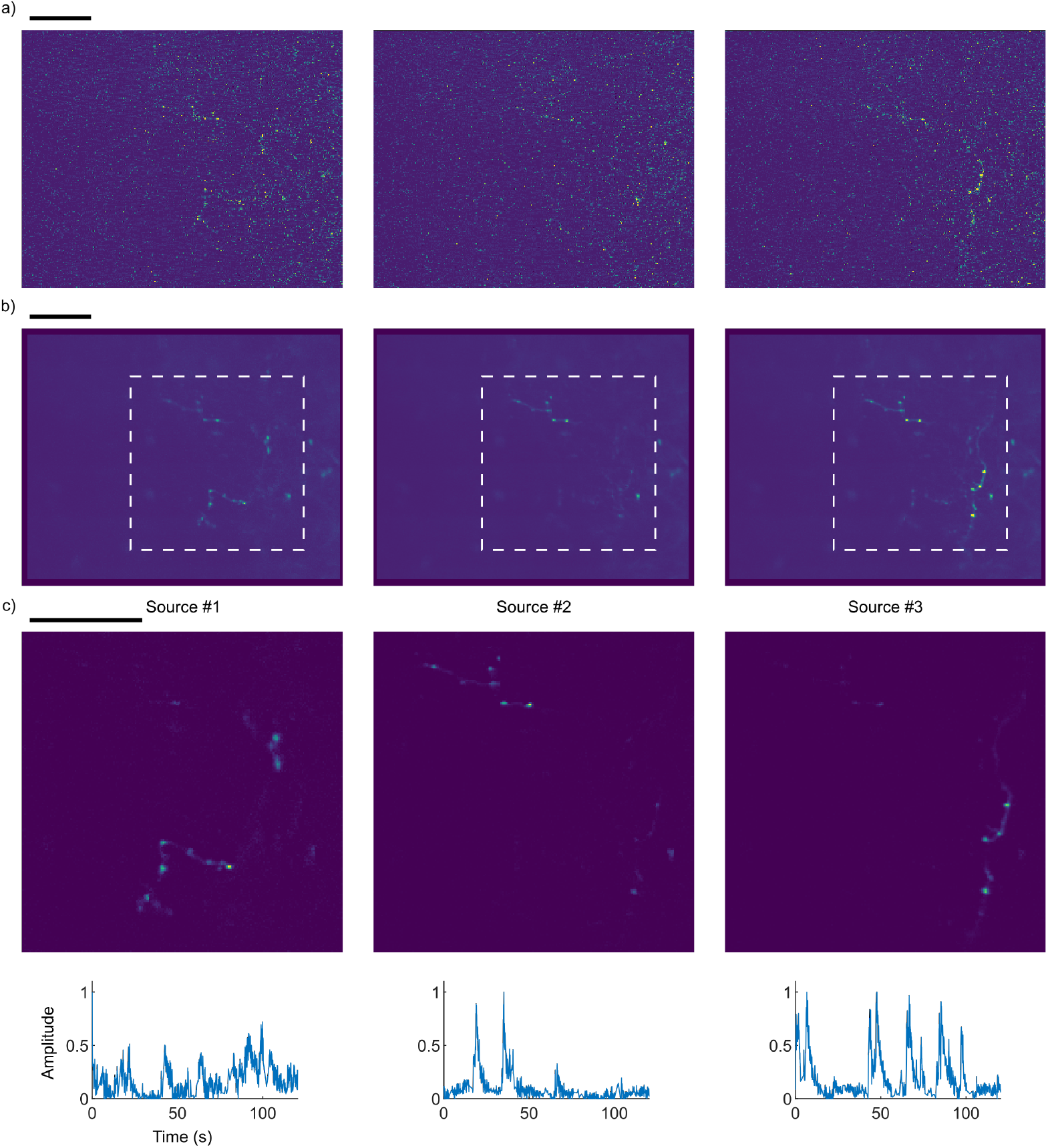
Extraction of axonal Ca^2+^ signals in in vivo microscopy image. a) Exemplar images of experiments of two-photon microscopy of Ca^2+^ in S1 in an awake mouse. b) Same image frames as (a) reconstructed as a noiseless spatio-temporal mixture of sources by NNeurite. c) Example of Ca^2+^ sources extracted by NNeurite for a 2 min image stack. Top: zoomed-in view of footprints for the dotted-white rectangular area in (a) and (b). Scale bars: 50 *μ*m.

## 5 Discussion

We have proposed novel methods for the analysis of microscopy image dedicated to *in vivo* temporal Ca^2+^ signals of small neurites: NNeurite. This complete solution takes unprocessed microscopy images and performs simultaneously image stabilization, denoising, spatial identification of an unknown number of Ca^2+^ sources in space and estimation of their temporal activity, all without human intervention. We have demonstrated the accuracy of NNeurite with sequences of images of axonal and dendritic features acquired in the behaving head-fixed mouse. Because methods able to automatically analyze data from *in vivo* microscopy of neurite activity are lacking, we provide NNeurite as a free and open-source Python library to fulfill this role and therefore help advancing our understanding of the micrometer-scale mechanisms supporting neuronal computation and cognition.

At the core of NNeurite is an original artificial NN which tackles the NMF problem for Ca^2+^ source separation. Our formulation markedly differs from standard methods for NMF-based Ca^2+^ image processing [5, 7] in that our training procedure learns an optimal network which has the ability to extract the amplitude of asynchronous Ca^2+^ sources, rather than optimizing directly the source amplitudes. It allows one to train the network with few chosen frames and reuse it at low computational cost for processing a large number of images – a feature which is particularly suited to long sequences of images of neurites with sparse activity in time. The overall complexity of NMF is tackled by considering smaller local problems on image patches. This strategy also allows, in principle, parallel computation on distributed architectures for large scale data processing [11].

The number of Ca^2+^ sources in the image sequence is automatically estimated by NNeurite, without human supervision. This is achieved at the level of the local NMF problems by sparsity-promoting group-lasso regularization [29] of both the temporal activity of sources and the footprints, and at the whole image level by non-supervised classification with an unknown number of clusters [31]. Importantly, our regularization strategy promotes the existence of only a few sources but this does not come at the cost of forcing the temporal activity or the spatial footprint of the identified sources to be sparse themselves, as would be the case with the more-popular *ℓ_1_* regularizer [5, 7, 9]. Beyond group-lasso regularization, NNeurite does not constrain further the shape and temporal activity of sources. This is crucial for neurite image processing where source shapes can take a complex, branched morphology. NNeurite was able to estimate high-resolution footprints with fine, micrometer-scale, morphological details such as dendritic spines and presynpatic boutons, hence demonstrating the benefits of this constraint-free approach. Another advantage of the flexibility of NNeurite is the inclusion in the same image model of other sources of fluorescence fluctuations which would do not conform well to a strict model of shape and amplitude, such as varying neuropil signals. It allows discarding their influence from the shapes and temporal amplitudes of the neurite sources. Posterior to NNeurite processing, an unsupervised, or supervised, classifier may be used to accurately discriminate sources of fluorescence of interest from others [11].

Imaging of micrometer-scale features in behaving animals is particularly sensitive to field-of-view non-rigid deformations due to small brain movements despite head-fixation. We took advantage of the flexibility of the proposed NN architecture to include non-rigid image stabilization as part of the NN-learning task. This is achieved by swapping the first image filtering layer of the NN with a 2D convolution layer (with no increase in the number of free NN parameters). A similar non-rigid Ca^2+^ image registration method based on local rigid transformations has been proposed before as a preprocessing for Ca^2+^microscopy images analysis [13]. The latter is particularly well-suited to images with strong stable, or slowly-changing, signals (as often seen with somatic fluorescence) which can be used as references to estimate local deformations with sub-pixel accuracy. Such reliable landmarks are not the rule with neurites imaging, most prominently for sparse axonal segments, because Ca^2+^ sources can be practically undetectable when inactive and background signals might not be as strong and distinctive. To solve this issue, we combined, for the first time, NMF-based source separation and image stabilization: images are registered to the footprints of the sources, taking into account their current amplitude, so that they serve as transient references in place of stable landmarks. While considering only integer spatial shifts, NNeurite was able to build high-resolution footprints for axons and dendrites and no duplicated sources were observed because of unaccounted image motion, hence demonstrating the accuracy of the proposed approach.

The proposed NN-based strategy is fundamentally different from NNs used for supervised soma segmentation in Ca^2+^ microscopy [6, 36–39] in that it is used in an unsupervised learning paradigm, much like autoencoders [27]. This makes the NN learning task into a regression-type problem which uses a NMF generative model, rather than a classification problem. To the best of our knowledge, it is the first use of unsupervised NNs for source-separation in neuronal Ca^2+^ microscopy. Supervised learning-based segmentation is a powerful strategy when large data sets labeled by humans are available and when the image properties and the features of the segmented objects are sufficiently similar from one experiment to another. To this day, challenging *in vivo* neurite imaging is not as widespread as somatic imaging and there are no large reference data sets with human annotations which can be used for supervised learning. Building such consensual data sets for *in vivo* neurite microscopy is complicated by the varied, unpredictable and complex shapes of neurites, unlike the more reproducible shapes of neuronal cell-bodies at low-resolution. We therefore argue for an unsupervised learning approach until varied and labeled reference data sets are available for *in vivo* neurite imaging.

NNeurite is a flexible framework for fluorescent image processing which we will enhance in the future. Implementing some forms of temporal regularization of Ca^2+^ traces [7] could be achieved with recurrent neuronal layers or grouped-images processing. Little adaption would be required to make it compatible with 3D+T image processing as the number of spatial dimensions are only accounted for by the first convolution layer. Furthermore, we have validated NNeurite use with neuronal Ca^2+^ images, but few aspects of the proposed methods are specific to Ca^2+^ signals. It therefore remains to be tested how well they perform with fluorescent sensors for other compounds [40, 41]) and membranar voltage [42, 43] in neurons.

## 6 Material and methods

### 6.1 Implementation details

NNeurite is implemented as a set of Python routines which use is demonstrated in Jupyter notebooks (https://github.com/nicolasC/NNeurite). Numpy and Scipy libraries are mainly used for mathematical computation. The proposed artificial NN structures are implemented thanks to the TensorFlow (2.0.0) dedicated library with GPU support.

### 6.2 *In vivo* Ca^2+^ imaging

C57Bl6/J mice were injected with a virus for GCaMP7f [44] expression in the primary whisking-related cortex (S1) at 400 *μ*m depth (L2/3 layer) and a glass window was implanted over S1 in place of the skull bone to allow microscopy imaging [45]. Three weeks after surgery, animals were habituated to head fixation and Ca^2+^ imaging was performed using a FemtoSmart two-photon laser scanning microscope (Femtonics, Budapest, Hungary) equipped with a 16x objective (0.8 NA, Nikon) (Fig 4a). Whiskers contralateral to the cranial window were intermittently stimulated using a glass-capillary attached to a piezoelectric actuator (PL-140.11 bender controlled by an E-650 driver; Physik Instrumente). A Ti:sapphire laser operating at *λ*=910 nm (Mai Tai DeepSee, Spectra-Physics) with an average excitation power at the focal point lower than 50 mW was used. The MES Software (MES v.4.6; Femtonics, Budapest, Hungary) was controlled the acquisition of images at 37 Hz with 0.635 *μ*m per pixel resolution. The size of the field of view was 325 *μ*m×271 *μ*m.

Animal experimental protocols were approved by the institutional ethical committee guidelines for animal research (Comité d’éthique de Bordeaux, no. 50DIR 15-A) and by the French Ministry of Research (agreement no.18892).

### 6.3 NNeurite processing parameters

All the analysis experiments relied on the same setup and parameters for NNeurite, except for an adapted regularization strength.

#### 6.3.1 Preprocessing

Out-of-focus images were first discarded by detecting global drops of fluorescence larger than 10% of the median fluorescence over a surrounding time window of 1.6 s. Resampling to 12.3 Hz (by a factor 3) was achieved by averaging subsequent frames and pixel values were homogeneously rescaled such as the minimum value was 0 and the 95% quantile was 0.1. The stack of images of dendrites used in Fig 5 was cropped to discard a large empty area. Images were then divided in patches of size 65 × 65 pixels.

#### 6.3.2 Artificial NNs

For local NNs (Fig 1), we used four fully connected layers for image analysis. The output of the second one (the ‘source layer’) computes the amplitude of the sources *q*^(0)^(*t*). We fixed a maximum of *K*_0_ = 8 sources per patch and used *K*_1_ = 16, *K*_2_ = 32 and *K*_2_ = 64. For regularization, we used *λ_M_* = 0.05 · var(*Y*), *λS* = 0.0005 · var(*Y*) for axonal image processing, and *λ_M_* = 0.01 var(*Y*), *λ_S_* = 0.0001 var(*Y*) for dendrites images. Soft non-negative constraints were used for the kernel weights of the final layer, which corresponds to image synthesis with *M* and *b*. Regularization of *M* was implemented as a group-lasso penalty for those weights. Regularization of *S* was implemented as a group-lasso penalty on batches of activities *q*^(0)^(*t*) of the source layer. When embedded image registration was used (Fig 3), larger overlapping image tiles were used so that the valid part of the convolution at the first analysis layer (2D convolution layer with unit strides and kernel size 65 × 65) was still of size 65 × 65 pixels. This architecture resulted in a total of around 311 000 free parameters to optimize for each image patch. We considered a maximum displacement of 5 pixels along each axis, and in each direction, hence resulting in 121 possible image shifts, or NN channels. For NN training, we ran 500 epochs of the Nadam SGD algorithm [46] with learning rate 0.005, a batch size of 32 and random shuffling of image frames between epochs.

#### 6.3.3 Local sources merging by affinity propagation clustering and refinement

We adapted to Python the code of Frey and Dueck for non-supervised clustering [31]. The affinity between local sources was defined as the Pearson correlation coefficient between their temporal amplitudes over the whole image sequence and the source preference was set to 0. A maximum of 5000 iterations were performed unless convergence was detected earlier. Source refinement was used with *k_max_* = 10 iterations of the alternated optimization procedure.

### 6.4 Computing architecture

All analysis experiments were performed using a Microsoft^®^ Windows^®^ 10 workstation equipped with an Intel^®^ Xeon^®^ CPU (2 processors with 6 cores each, 3.40 GHz maximum), a NVIDIA^®^ GeForce^®^ RTX 2080 Ti GPU (4352 CUDA cores, 544 Tensor cores, 1350 MHz base clock) and 64 GB RAM (2400 MHz). The tensorflow library for NN optimization was set up to take advantage of GPU parallel processing with the cuDNN library.

## 7 Funding

The authors declare no competing financial interests. NC received funding from the Marie Sklodowska-Curie individual fellowship under the European Union’s Horizon 2020 research and innovation program (AXO-MATH, grant agreement n^®^ 798326). F.G. received funding from the Agence Nationale de la Recherche (SyTune, ANR-21-CE37-0010), the European Research Council under the European Union’s Horizon 2020 research and innovation program (NEUROGOAL, grant agreement n^®^ 677878), the Region Nouvelle-Aquitaine and the University of Bordeaux. RWT was supported by research grants from the National Institute of General Medical Sciences (NIGMS, GM058234); National Institute of Neurological Disorders and Stroke (NINDS, U19NS107616); the National Institute of Mental Health (NIMH, MH071739); and the Simons, Mathers, and Burnett Family Foundations.

## Footnotes

1 https://github.com/nicolasC/NNeurite

## Notes

### Competing Interest Statement

The authors have declared no competing interest.

